# The Rab11 effectors Fip5 and Fip1 regulate zebrafish intestinal development

**DOI:** 10.1101/2020.06.15.153569

**Authors:** Cayla E. Jewett, Bruce H. Appel, Rytis Prekeris

**Affiliations:** Department of Cell and Developmental Biology, University of Colorado School of Medicine, Aurora, Colorado 80045 USA; Department of Pediatrics, University of Colorado School of Medicine, Aurora, Colorado 80045 USA

**Keywords:** FIP5, FIP1, Rip11, RCP, Rab11, microvilli, keratin, Rab7, MVID

## Abstract

The Rab11 apical recycling endosome pathway is a well-established regulator of polarity and lumen formation; however, Rab11-vesicular trafficking also directs a diverse array of other cellular processes, raising the question of how Rab11 vesicles achieve specificity in space, time, and content of cargo delivery. In part, this specificity is achieved through effector proteins, yet the role of Rab11 effector proteins *in vivo* remains vague. Here, we use CRISPR/Cas9 gene editing to study the role of the Rab11 effector Fip5 during zebrafish intestinal development. Zebrafish contain two paralogous genes, *fip5a* and *fip5b*, that are orthologs of human *FIP5*. We find that *fip5a* and *fip5b* mutant fish show phenotypes characteristic of microvillus inclusion disease, including microvilli defects, inclusion bodies, and lysosomal accumulation. Single and double mutant analysis suggest that *fip5a* and *fip5b* function in parallel and regulate apical trafficking pathways required for assembly of keratin at the terminal web. Remarkably, in some genetic backgrounds, the absence of Fip5 triggers protein upregulation of a closely related family member, Fip1. This compensation mechanism occurs both during zebrafish intestinal development and in tissue culture models of lumenogenesis. In conclusion, our data implicate the Rab11 effectors Fip5 and Fip1 in a trafficking pathway required for apical microvilli formation.

## INTRODUCTION

Development of many organs, such as the gastrointestinal system, kidneys, and respiratory tract requires morphogenetic remodeling of cells to form a hollow tube, or lumen (Jewett and Prekeris, 2018). Whereas the mechanisms cells use to form a lumen vary by organ, a common feature is that cells adopt a highly polarized conformation including establishment of apical structures such as primary cilia, motile cilia, or microvilli (Apodaca and Gallo, 2013). Intestinal epithelia are one of the few vertebrate cell types to lack primary cilia, but their apical cell surface is covered with a brush border composed of actin-rich membrane protrusions called microvilli to aid in nutrient absorption (Apodaca and Gallo, 2013). The molecular basis of cell polarization is well defined, but much less is understood about how trafficking pathways govern formation of these apical structures, especially *in vivo.*

The Rab11 apical recycling endosome pathway is a well-established regulator of polarity and lumen formation (Jewett and Prekeris, 2018). However, Rab11-directed trafficking events are also implicated in a number of other cellular processes, raising the question of how Rab11 vesicles achieve specificity when involved in numerous cellular functions. In part, Rab specificity is achieved through interaction with effector proteins, and Rab11 in particular interacts with a family of effector proteins called Rab11-Family Interacting Proteins (FIPs) (Horgan and McCaffrey, 2009). There are five FIP family members, all of which contain a coiled-coil region at the C-terminus of the protein, allowing dimerization and binding to two Rab11 molecules, effectively forming a functional heterotetramer. Different FIPs appear to function in unique cellular processes including cytokinesis (FIP3 and FIP4), ciliogenesis (FIP3), and cargo recycling to the cell surface (FIP1, FIP2, FIP5) (Horgan and McCaffrey, 2009). Previously, our lab implicated FIP5 in apical lumen formation in 3D Madin Darby Canine Kidney (MDCK) cell culture. We and others have shown that Rab11-FIP5 endosomes are required for lumenogenesis and interact with the actin binding protein MYO5B to traffic cargo to the apical cell surface (Lapierre et al., 2001, Willenborg et al., 2011, Mangan et al., 2016). However, whether FIP5 plays a role in coordinating lumen morphogenesis during development *in vivo* is unknown.

Polarization is critical for cell function such that polarity disruption results in a number of diseases. Microvillus Inclusion Disease (MVID) is one such example, arising from the inability to form and maintain microvilli at the apical cell surface (Al-Daraji et al., 2010). Patients with MVID suffer from intractable diarrhea and malabsorption due to absent or very sparse microvilli and typically do not live past childhood. At the cellular level, patients display characteristic trafficking defects of lysosome accumulation and inclusion bodies containing microvilli (Phillips et al., 1985, Phillips and Schmitz, 1992, Ruemmele et al., 2006). Mutations in MYO5B are found in patients with MVID and mutations in the zebrafish ortholog *myoVb* (also called *goosepimples*), result in inclusion bodies and trafficking defects (Müller et al., 2008, Ruemmele et al., 2010, Sidhaye et al., 2016). Moreover, experiments from intestinal tissue culture models suggest that the interaction between Rab11 and MYO5B is essential for microvilli maintenance (Knowles et al., 2014). Given that FIP5 interacts with MYO5B and is required for lumen formation in tissue culture, we hypothesized that FIP5 regulates intestinal development and microvilli formation *in vivo*.

## RESULTS AND DISCUSSION

The mechanisms by which cells polarize and form an apical lumen have been studied extensively in 3D tissue culture, but whether these processes are recapitulated *in vivo* is unclear, because vertebrate models are inherently more complex and have compensatory mechanisms. Furthermore, intestinal tissue culture models are limited due to a lack of proper microvilli that are subject to the stresses and strains encountered by a functional animal intestine. To address these limitations, we utilized zebrafish intestinal development as an *in vivo* model of lumenogenesis and microvilli formation. We first examined the degree to which zebrafish Fip5 protein was conserved with human and dog FIP5 protein, as most work on FIP5 during cell polarization has been performed in MDCK cells. Zebrafish contain Fip5a and Fip5b orthologs to mammalian FIP5 with two highly conserved functional domains: a phospholipid-binding domain C2 domain at the N-terminus and a coiled-coil region at the C-terminus of the protein (Figure S1A, yellow and blue highlight, respectively) required for dimerization and binding to Rab11 (Prekeris et al., 2001). Zebrafish intestinal development begins around 3 days post-fertilization (dpf) when many small lumens develop throughout the intestinal tract and subsequently fuse to form a single continuous lumen from mouth to anus (Ng et al., 2005, Alvers et al., 2014). To determine where *fip5a* and *fip5b* were expressed in zebrafish larvae during development, we performed *in situ* hybridization on 4 dpf larvae. Luminal organs such as the intestine, spinal cord, and notochord expressed *fip5a* and *fip5b* (Figure S2A, B). In measuring mRNA levels of *fip5a* and *fip5b*, we found that both transcripts showed increased levels around 3 dpf, and high levels of *fip5b* mRNA persisted throughout 8 dpf (Figure S2C). We thus focused our efforts first on *fip5b*.

### Endosome maturation and terminal web keratin organization require Fip5b function

To study the function of Fip5b, we used CRISPR/Cas9 gene editing. We selected two different *fip5b* alleles that introduced a premature stop codon right after the C2 domain at the N-terminus (Figure 1A, Figure S1B), thereby eliminating the Rab-binding domain (RBD) at the C-terminus essential for Fip5 function. We maintained these *fip5b* mutant stocks in a heterozygous state and performed intercrosses to generate zygotic mutants for analysis. Stage matched wild-type siblings were used as controls. We performed qRT-PCR to measure *fip5b* expression in *fip5b^CO40^* homozygous mutant larvae and observed an almost complete loss of *fip5b* mRNA levels (Figure 1B), suggestive of nonsense-mediated decay. *fip5b^CO40^* homozygous mutant fish appeared morphologically normal from embryo through adulthood and were homozygous viable as adults. However, to determine if loss of Fip5b affected intestinal development at the cellular level, we performed transmission electron microscopy on fixed sections through the midgut region (Figure 1C, yellow box) at developmental time points. At 3 dpf, when intestinal lumen morphogenesis initiated, *fip5b^CO40^* mutant larvae formed a single lumen (Figure 1D), but upon closer examination, we noticed an accumulation of membrane vesicles in the subapical cytoplasm not present in wild type larvae (Figure 1E yellow box, F). These vesicles resembled inclusion bodies which are pathological hallmarks of MVID. At 6 dpf when intestinal development was mostly complete (Ng et al., 2005), inclusion-like bodies were no longer evident near the subapical surface, consistent with MYO5B mutant mice in which microvillus inclusions were more pronounced in neonates and disappeared after weaning due to decreased apical macropinocytosis (Knowles et al., 2014, Weis et al., 2016). Instead, intestinal cells of homozygous mutant larvae showed an accumulation of small (less than 500nm) apical vesicles (Figure 1G, H) and large (greater than 500nm) organelles that resided medially in the cells (Figure 1G arrows, I) compared to wild-type cells which did not show an accumulation of intracellular vesicles. Moreover, microvilli were shorter in both the anterior intestinal bulb and posterior midgut of 6 dpf homozygous mutant fish compared to wild-type siblings (Figure 1G, J). Finally, the terminal web, an apical cytoskeletal network anchoring microvilli into the cell, was disrupted in mutant fish. Wild-type larvae had a defined electron dense line at the base of the microvilli and an organelle-free zone just below the apical cell surface which was absent in mutants (Figure 1G, brackets). These data revealed trafficking and microvilli defects in *fip5b^CO40^* mutant larvae.

**Figure 1.**
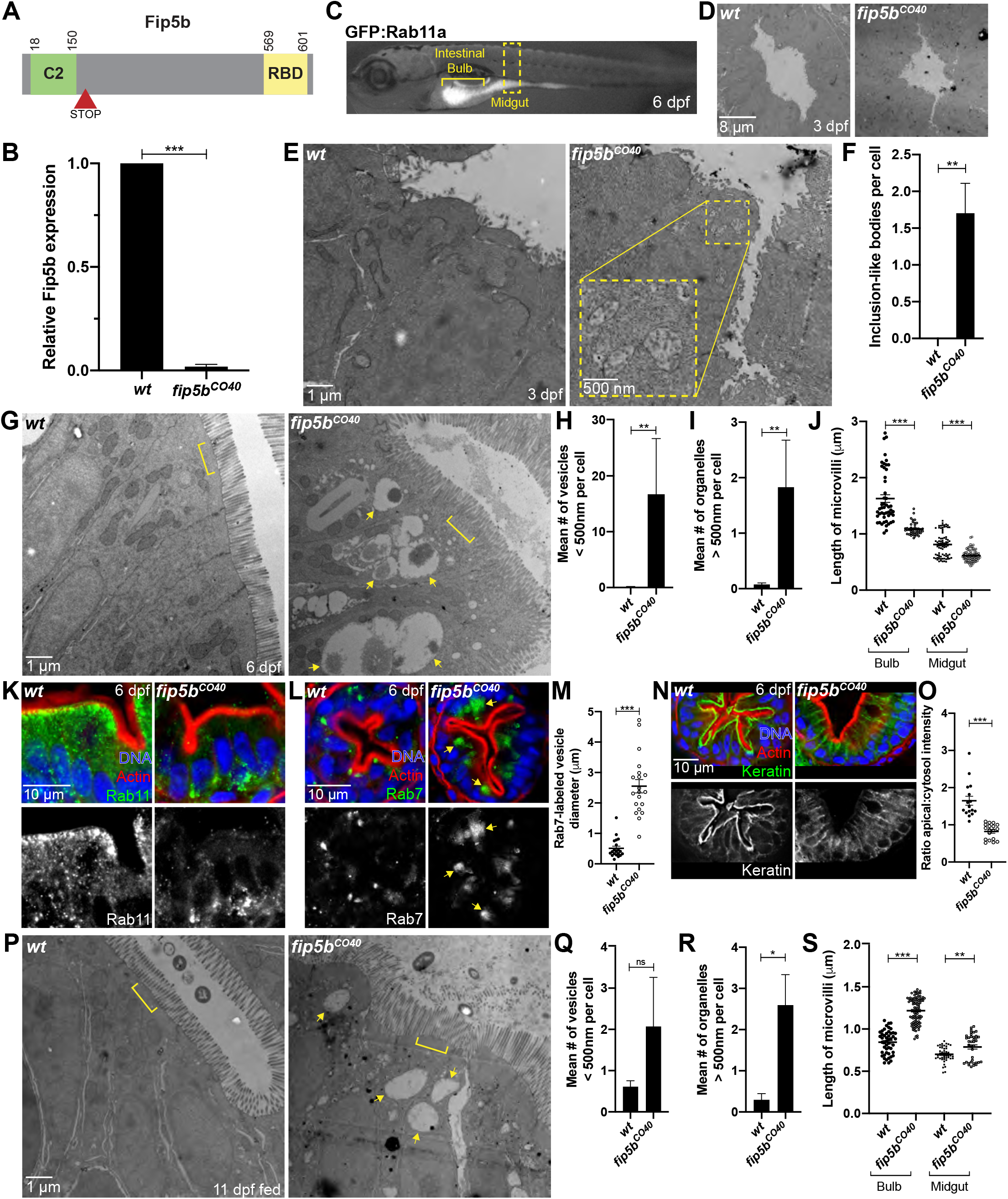
Endosome maturation and terminal web keratin organization require Fip5b function. (A) Domain schematic of zebrafish Fip5b protein containing a C2 domain at the N-terminus and a Rab-binding domain (RBD) at the C-terminus. The red arrowhead with STOP denotes premature termination codon in *fip5b* mutant alleles. (B) qRT-PCR for *fip5b* in wild-type and *fip5b^CO40^* mutant larvae at 6 dpf. (C) 6 dpf larvae expressing *Tg(hsp:GFP:Rab11a)* labeling the intestine. The intestinal bulb is denoted by a bracket and the midgut by a dashed box. All following images are representative cross sections through the midgut region. Wild-type siblings are used as controls. (D) Electron micrographs showing 3 dpf wild-type and *fip5b^CO40^* mutant larvae. Luminal space is lighter gray region. (E) High magnification electron micrographs showing 3 dpf wild-type and *fip5b^CO40^* mutant larvae. Yellow box shows zoomed in view on a region with subapical inclusion-like bodies. (F) Quantitation of the mean number of inclusion-like bodies per cell in 3 dpf wild-type and *fip5b^CO40^* mutant larvae. (G) Electron micrographs showing 6 dpf wild-type and *fip5b^CO40^* mutant larvae. Arrows point to larger than 500nm organelles and brackets mark terminal web or lack thereof in mutants. (H) Quantitation of less than 500nm apical vesicles in 6 dpf larvae. (I) Quantitation of greater than 500nm organelles in 6 dpf larvae. (J) Quantitation of microvilli length in the intestinal bulb and midgut in 6 dpf larvae. (K,L,N) Immunohistochemistry on cross sections of 6 dpf wild-type and *Fip5b^CO40^* mutant larvae stained with Hoechst (blue), Phalloidin (red), and Rab11 (K), Rab7 (L), or cytokeratin (N) (green). (M) Quantitation of Rab7-vesicle diameter. (O) Ratio of fluorescence intensity of apical keratin to cytoplasmic keratin. (P) Electron micrographs showing 11 dpf fed wild-type and *fip5b^CO40^* mutant larvae. Arrows point to larger than 500nm organelles and brackets mark terminal web or lack thereof in mutants. (Q) Quantitation of less than 500nm apical vesicles in 11 dpf larvae. (R) Quantitation of greater than 500nm organelles in 11 dpf larvae. (S) Quantitation of microvilli length in the intestinal bulb and midgut of 11 dpf larvae. All plots show mean with standard error of the mean. A t-test was used for Gaussian data and a Mann-Whitney test for all other statistics. ***P < 0.0005, **P < 0.005, *P < 0.05.

To investigate the identity of the large organelles observed in *fip5b^CO40^* mutant larvae intestinal cells, we performed immunohistochemistry to detect proteins that serve as common endosome markers. Because Fip5 binds Rab11 vesicles, we first examined Rab11 localization. In wild-type intestinal cells, Rab11 vesicles localized just beneath the apical cell surface, as revealed by actin staining (Figure 1K). In contrast, Rab11 vesicles mislocalized to the basolateral surface of intestinal cells in *fip5b^CO40^* mutant larvae (Figure 1K). Because the large organelles observed through electron microscopy in mutant tissue were near the apical cell surface, they were unlikely to be Rab11-positive. Intestinal cells of MVID patients accumulate lysosomal granules (Iancu et al., 2007), so we next stained cells to detect the late endosome/lysosome marker Rab7. Notably, Rab7-positive organelles accumulated near the apical cell surface in *fip5b^CO40^* mutant cells, whereas we did not detect these large organelles in wild-type cells (Figure 1L). These Rab7 endosomes were consistent in size and localization with the structures revealed by electron microscopy (Figure 1M). Taken together, these data suggested that Fip5b is required for Rab11 apical localization and Rab7 endosomal trafficking processes.

Our electron microscopy analysis also revealed defects in microvilli length and the terminal web in *fip5b^CO40^* mutant cells. The terminal web is composed of actin and intermediate filaments and is located just below the apical cell surface to anchor the base of microvilli into the cell (Mooseker et al., 1984). Because actin localized to the apical cell surface of mutant cells similar to wild-type cells (Figure 1K, L, N), we focused our attention on intermediate filaments. In polarized epithelia, intermediate filaments are composed of keratin polymers, so we stained cells with a pan-cytokeratin antibody to visualize intermediate filaments comprising the terminal web. We found that in wild-type cells, the keratin network resided just below the apical actin network; however, in *fip5b^CO40^* mutant cells, keratin mislocalized to lateral and cytoplasmic regions of the cell (Figure 1N, O). These observations were consistent with the possibility that Fip5b regulates keratin polymerization and terminal web formation at the apical cell surface.

Terminal web defects result in microvilli abnormalities, which can be exacerbated by physical stress from intestinal activity. We therefore hypothesized that fed mutant larvae would show more severe microvilli phenotypes than unfed 6 dpf larvae still living off the yolk. To test this, we began feeding the larvae daily at 7 dpf and then analyzed larvae at 11 dpf. Mutant larvae showed moderate trafficking defects at 11 dpf (Figure 1P, arrows, Q, R); however, the terminal web defects recovered, and microvilli were now significantly longer than wild-type siblings (Figure 1P, bracket, S). This phenotypic recovery was unexpected and perhaps explains in part why adult mutant fish were homozygous viable. Importantly, these trafficking and microvilli phenotypes were recapitulated in another *fip5b* mutant allele *fip5b^CO43^* (Figure S1B, S3A-D) indicating that these phenotypes were specific to *fip5b*. Taken together, these data provided evidence that Fip5b functions in apical trafficking processes and microvilli formation during zebrafish intestinal development.

### fip5a functions similarly to fib5b in endosome maturation and terminal web organization

Whereas f*ip5b* mutant phenotypes were prominent during early developmental stages, these mutant fish recovered from these defects and were viable as homozygous adults. One possible explanation is a compensatory mechanism, perhaps through upregulation of another trafficking pathway, and an obvious candidate for compensation is the zebrafish *fip5b* paralog, *fip5a*. To test Fip5a’s role in intestinal development, we again used CRISPR to create *fip5a* mutant alleles (Figure 2A, Figure S1C). *fip5a* mutant stocks were maintained in a heterozygous state and intercrossed to generate zygotic mutants for analysis. Stage matched wild-type siblings were used as controls. Similar to *fip5b* mutants, *fip5a^CO38^* homozygous mutant larvae were morphologically normal and viable as homozygous adults. To study the role of *fip5a* during intestinal development, we performed the same transmission electron microscopy analysis on fixed sections through the mid-intestinal region. Notably, *fip5a^CO38^* mutant fish recapitulated phenotypes seen in *fip5b* mutant fish. At 3 dpf, *fip5a^CO38^* mutant larvae formed a lumen, but exhibited subapical organelles resembling inclusion bodies (Figure 1B, C). By 6 dpf, inclusion bodies cleared, and *fip5a^CO38^* mutant cells now accumulated small apical vesicles (Figure 1D, E) and large organelles (Figure 1D arrows, F) not present in wild-type larvae. Additionally, midgut microvilli were shorter (Figure 1D, G) and the terminal web was also disrupted in mutants compared to wild-type larvae (Figure 1D, brackets). These large organelles were Rab7-positive in *fip5a^CO38^* mutant fish and terminal web defects appeared to be the result of mislocalized keratin from the apical cell surface (Figure 2H-K). Again, similar to *fip5b* mutants at 11 dpf, *fip5a^CO38^* mutants maintained trafficking defects (Figure 1L, arrows, M, N), but unlike *fip5b* mutants, the terminal web defects and shorter microvilli persisted in *fip5a^CO38^* mutants at 11 dpf (Figure 1L, brackets, O). Importantly, these trafficking and microvilli phenotypes were recapitulated in another *fip5a* mutant allele, *fip5a^CO35^* (Figure S1C, S3E-H) indicating that these phenotypes were specific to *fip5a*. Collectively, these data implicated Fip5a in apical trafficking and microvilli formation and suggested a similar function to Fip5b during zebrafish intestinal development.

**Figure 2.**
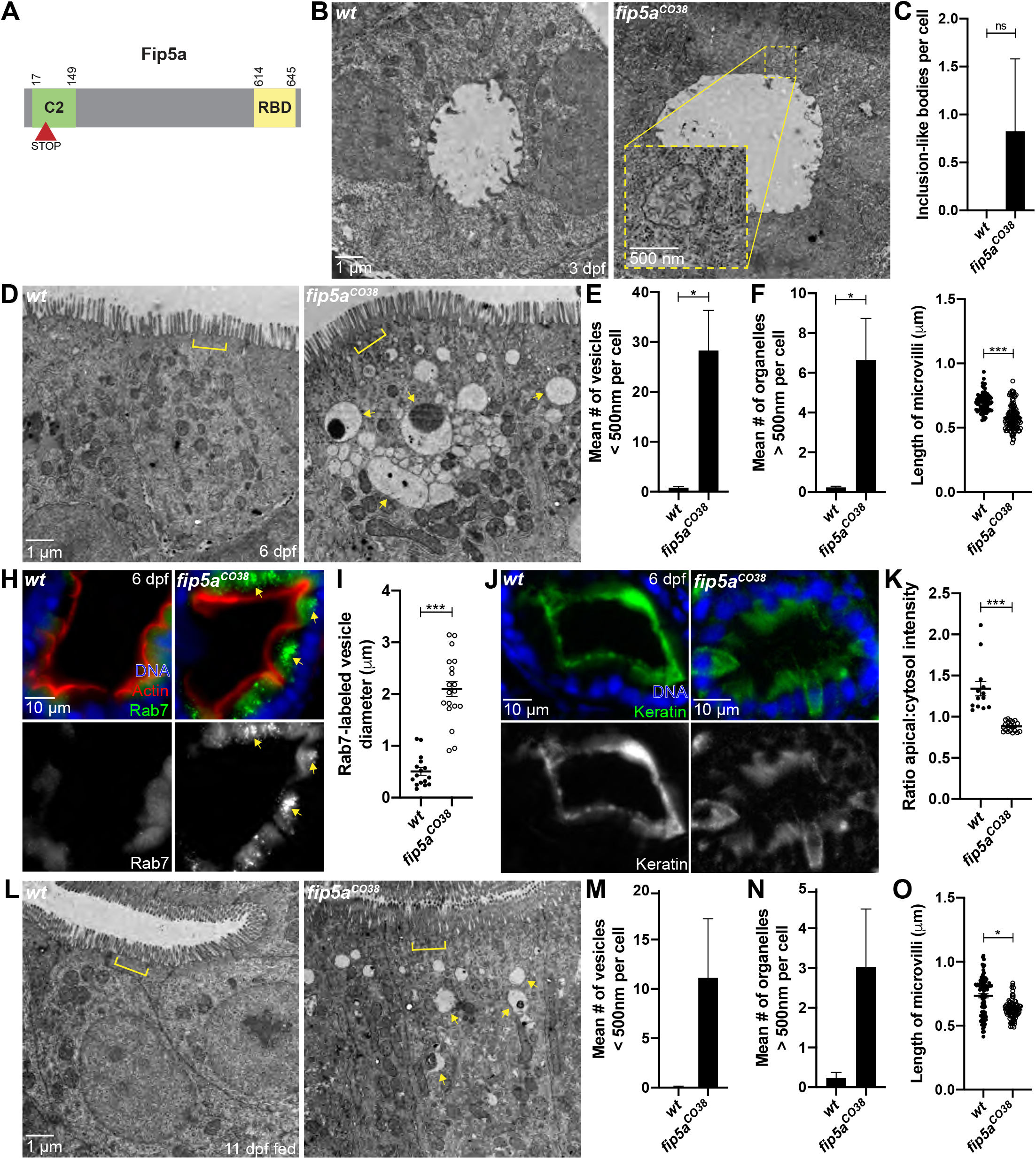
*fip5a* functions similarly to *fib5b* in endosome maturation and terminal web organization. (A) Domain schematic of zebrafish Fip5a protein containing a C2 domain at the N-terminus and a Rab-binding domain (RBD) at the C-terminus. The red arrowhead with STOP denotes premature termination codon in *fip5a* mutant alleles. All following images are representative cross sections through midgut region. Wild-type siblings are used as controls. (B) Electron micrographs showing 3 dpf wild-type and *fip5a^CO38^* mutant larvae. Yellow box shows zoomed in view on a region with subapical inclusion-like bodies. (C) Quantitation of the mean number of inclusion-like bodies per cell in 3 dpf wild-type and *fip5a^CO38^* mutant larvae. (D) Electron micrographs showing 6 dpf wild-type and *fip5a^CO38^* mutant larvae. Arrows point to larger than 500nm organelles and brackets mark terminal web or lack thereof in mutants. (E) Quantitation of less than 500nm apical vesicles in 6 dpf larvae. (F) Quantitation of greater than 500nm organelles in 6 dpf larvae. (G) Quantitation midgut microvilli length in 6 dpf larvae. (H) Immunohistochemistry on cross sections of 6 dpf wild-type and *fip5a^CO38^* mutant larvae stained with Hoechst (blue), Phalloidin (red), and Rab7 (green). (I) Quantitation of Rab7-vesicle diameter. (J) Immunohistochemistry on cross sections of 6 dpf wild-type and *fip5a^CO38^* mutant larvae stained with Hoechst (blue) and cytokeratin (green). (K) Ratio of fluorescence intensity of apical keratin to cytoplasmic keratin. (L) Electron micrographs showing 11 dpf fed wild-type and *fip5a^CO38^* mutant larvae. Arrows point to larger than 500nm organelles and brackets mark terminal web or lack thereof in mutants. (M) Quantitation of less than 500nm apical vesicles in 11 dpf larvae. (N) Quantitation of greater than 500nm organelles in 11 dpf larvae. (O) Quantitation midgut microvilli length in 11 dpf larvae. All plots show mean with standard error of the mean. A t-test was used for Gaussian data and a Mann-Whitney test for all other statistics. ***P < 0.0005, *P < 0.05.

### fip5a and fip5b double mutants show severe microvilli and trafficking phenotypes

*fip5a* and *fip5b* homozygous mutant larvae showed similar phenotypes, but it remained unclear whether *fip5a* and *fip5b* function in parallel or through a common pathway. To test this, we created a *fip5a; fip5b* heterozygous mutant line (*fip5a^CO35/+^; fip5b^CO40/+^*). This fish line was maintained in a heterozygous state and intercrossed to generate *fip5a^CO35/CO35^; fip5b^CO40/CO40^* homozygous double mutant embryos for experiments. Wild-type siblings were used as controls. Through electron microscopy analysis at 6 dpf, *fip5a^CO35^; fip5b^CO40^* zygotic double mutant fish showed two classes of phenotypes. The first was a severe microvilli defect where microvilli density was significantly reduced and the microvilli that did form were shorter and more heterogeneous in double mutants compared to wild-type larvae (Figure 3A and A’’, braces, C). The second was a severe trafficking phenotype where the majority of the cell cytosol was filled with giant Rab7-positive organelles (Figure 3A’ and A’’’, arrows, D, E). Double mutant fish also accumulated small apical vesicles and terminal web defects (Figure 3A’’’, bracket, F, G) like those seen in single mutants. These phenotypes were not mutually exclusive, as some mutant larvae displayed both microvilli and trafficking defects. It is worth noting that wild-type siblings also showed mild microvilli, terminal web, and trafficking defects (Figure 3A-A’, brace, bracket, and arrows, respectively), perhaps suggestive of maternal contribution, as stage-matched wild-type AB fish did not show these phenotypes (Figure 3B). In addition to these intestinal phenotypes, about 50% of the double mutant larvae had multiple kidney lumens, whereas wild-type siblings or single *fip5a* or *fip5b* mutant larvae always had a single continuous kidney lumen (Figure 3H, arrows). Moreover, double mutant animals did not live past two weeks. Thus, the severity of these double mutant phenotypes suggested that Fip5a and Fip5b function in parallel in microvilli formation during zebrafish intestinal development through apical trafficking pathways that regulate terminal web formation.

**Figure 3.**
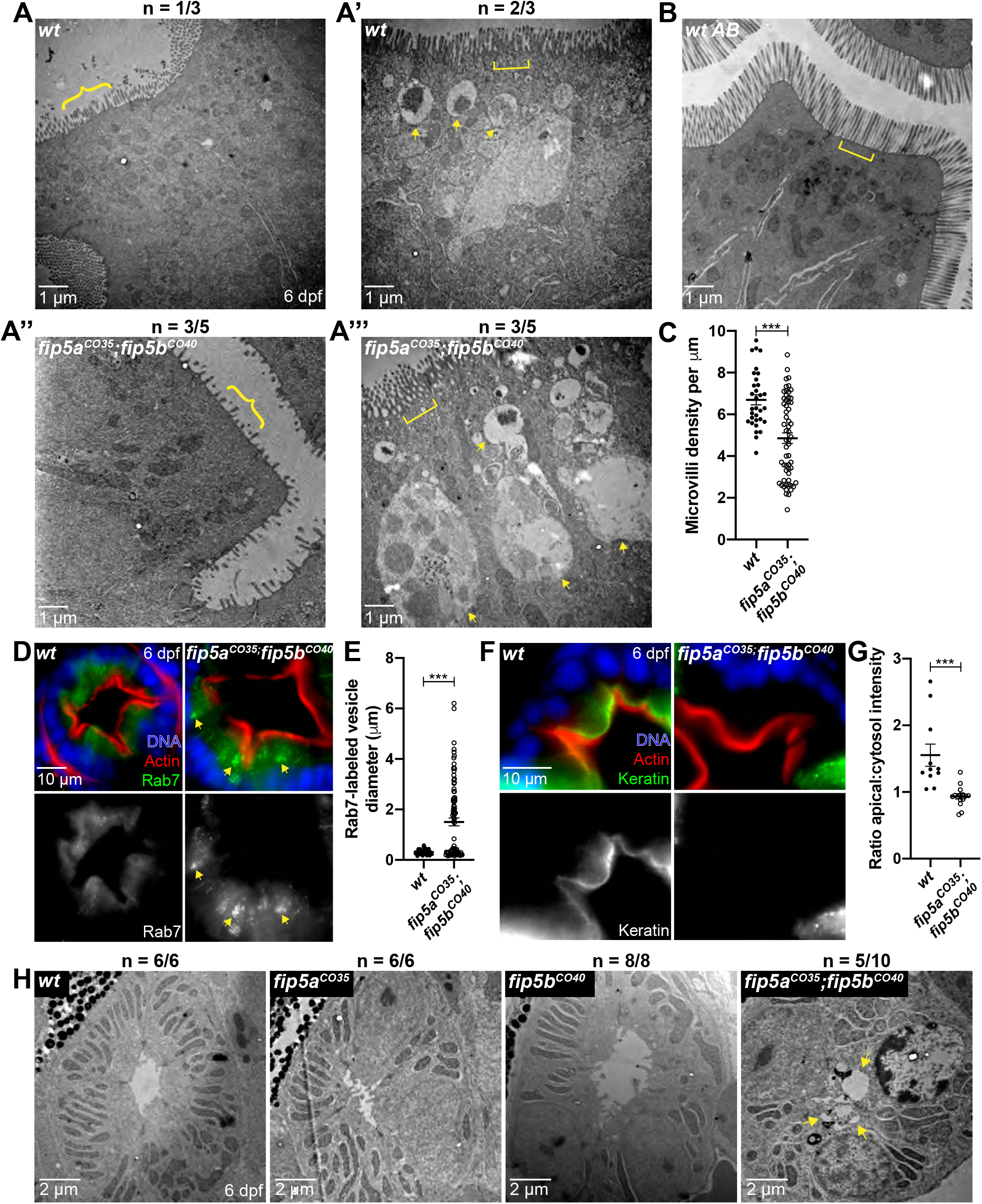
*fip5a* and *fip5b* double mutants show severe microvilli and trafficking phenotypes. All following images are representative cross sections through the midgut region of 6 dpf larvae. (A-A’’’’) Electron micrographs showing wild-type siblings and *fip5a^CO35/CO35^; fip5b^CO40/CO40^* zygotic mutant larvae. Arrows point to larger than 500nm organelles, braces point out sparse microvilli, and brackets mark terminal web or lack thereof in mutants. N indicates number of representative larvae out of total number of larvae analyzed. (B) Electron micrograph showing wild-type AB larva. (C) Quantitation of microvilli density. (D) Immunohistochemistry on cross sections of wild-type and *fip5a^CO35^; fip5b^CO40^* mutant larvae stained with Hoechst (blue), Phalloidin (red), and Rab7 (green). (E) Quantitation of Rab7-vesicle diameter. (F) Immunohistochemistry on cross sections of wild-type and *fip5a^CO35^; fip5b^CO40^* mutant larvae stained with Hoechst (blue), Phalloidin (red) and cytokeratin (green). (G) Ratio of fluorescence intensity of apical keratin to cytoplasmic keratin. (H) Electron micrographs of kidneys in wild-type, *fip5b^CO40^* mutant, *fip5a^CO35^* mutant, and *fip5a^CO35^; fip5b^CO40^* double mutant larvae. N indicates number of representative kidneys out of total number of kidneys analyzed. Arrows point to multiple lumens in *fip5a^CO35^; fip5b^CO40^* double mutant larvae. All plots show mean with standard error of the mean. A t-test was used for Gaussian data and a Mann-Whitney test for all other statistics. ***P < 0.0005.

### Upregulation of Fip1 rescues fip5a and fip5b double mutant phenotypes

Although larvae deficient for zygotic functions of both *fip5a* and *fip5b* had severe intestinal phenotypes, contribution of wild-type maternal products to the eggs laid by heterozygous females potentially partially suppressed the phenotype. To test this possibility, we removed the maternal contribution of *fip5a* by intercrossing *fip5a^CO35/CO35^; fip5b^CO40/+^* adults. We called these maternal-zygotic double mutants *fip5a^CO35^; fip5b^CO40^ mat-* to differentiate from zygotic double mutants in Figure 3 created from a heterozygous intercross (Figure 4A versus B). Surprisingly, *fip5a^CO35^; fip5b^CO40^ mat-* larvae, lacking maternal and zygotic functions of *fip5a* and zygotic functions of *fip5b*, had no intestinal phenotypes and could not be discerned morphologically from wild-type larvae (Figure 4C). Thus, removing maternal *fip5a* function suppressed, rather than enhanced, the phenotype of double mutant larvae.

**Figure 4.**
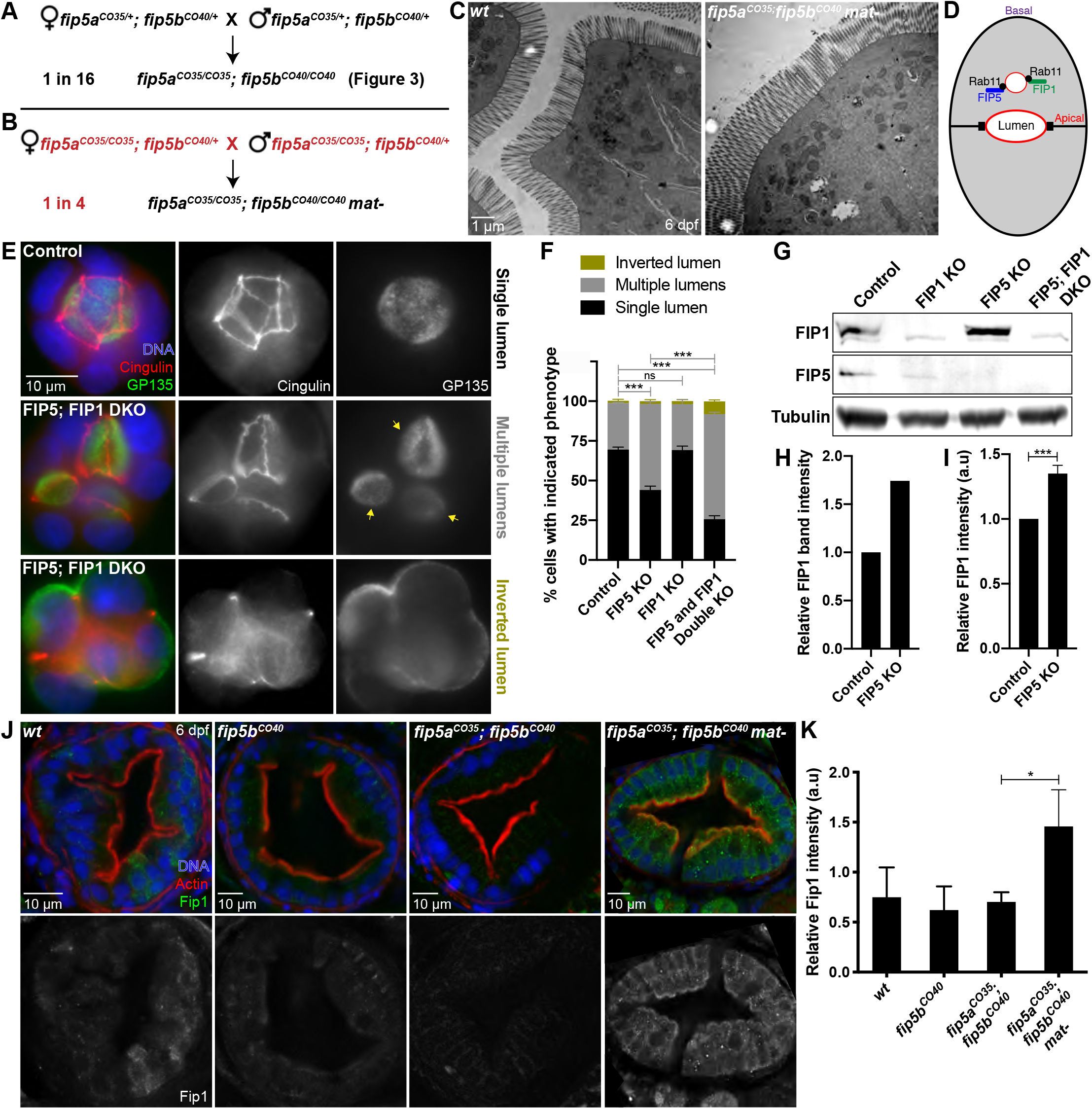
Upregulation of Fip1 rescues *fip5a* and *fip5b* double mutant phenotypes. (A-B) Schematic of genetic crosses resulting in *fip5a^CO35^; fip5b^CO40^* double zygotic or maternal/zygotic (mat-) mutant offspring generated from two different parental genotypes. (C) Electron micrographs showing cross sections through midgut of 6 dpf wild-type AB and *fip5a^CO35^; fip5b^CO40^ mat-* mutant larvae. (D) Cartoon schematic showing FIP5 and FIP1 bind same Rab11 vesicles. (E) Wild-type and FIP5; FIP1 double KO MDCK cells grown in an extracellular matrix to induce 3D lumen formation. Arrows denote multiple lumens in KO cyst. (F) Quantitation of luminal phenotypes. (G) Western Blot on wild-type and KO MDCK cell lysates probed for FIP1, FIP5 or tubulin (control) antibodies. (H) Quantitation of FIP1 band intensity for wild-type and FIP5 KO cell lysates. (I) Quantitation of FIP1 fluorescence intensity in wild-type and FIP5 KO cells grown in polarized monolayers. Representative images are shown in Figure S4A. (J) Immunohistochemistry on cross sections through midgut of 6 dpf wild-type, *fip5b^CO40^* mutant, *fip5a^CO35^; fip5b^CO40^* double mutant, and *fip5a^CO35^; fip5b^CO40^ mat-* mutant larvae stained with Hoechst (blue), Phalloidin (red) and Fip1 (green). (K) Quantitation of fluorescence intensity of Fip1. All plots show mean with standard error of the mean. A t-test was used for Gaussian data and a Mann-Whitney test for all other statistics. ***P < 0.0005, *P < 0.05.

Recent literature has posited a role for compensatory mechanisms due to gene knockout when the mutant mRNA undergoes nonsense-mediated decay (Rossi et al., 2015, El-Brolosy et al., 2019). One compensatory mechanism included upregulation of transcripts similar in sequence to the mRNA encoded by the mutated gene (El-Brolosy et al., 2019). We thus wondered if another Fip family member could be upregulated in the absence of maternal and zygotic functions of *fip5a* and zygotic functions of *fip5b*.

Previous work in our lab showed that both FIP5 and FIP1 bind the same Rab11 vesicles and FIP5 proteomics revealed an interaction with FIP1 (Willenborg et al., 2011, Mangan et al., 2016) (Figure 4D). We thus used a MDCK tissue culture model of lumenogenesis to ask if FIP1 could compensate for FIP5. When MDCK cells were grown in an extracellular matrix, the majority of wild-type cells formed a single continuous lumen inside the cyst of cells; however, most FIP5 and FIP1 double KO cells showed a multilumenal phenotype and a small percentage showed an inverted polarity phenotype (Figure 4E, F). These luminal phenotypes were significantly more severe than FIP5 KO alone (Figure 4F). Correspondingly, Western Blot analysis demonstrated that FIP1 protein levels were upregulated in FIP5 KO cells (Figure 4G, H) and immunohistochemistry experiments with a FIP1 antibody confirmed this (Figure 4I, Figure S4A). This protein upregulation was specific to FIP1 in FIP5 KO cells, as FIP5 levels did not increase in FIP1 KO cells (Figure 4G, Figure S4B). Moreover, FIP5 and FIP1 double KO cells did not show general defects in apical polarity or tight junction formation when grown in a polarized monolayer (Figure S4C, D), suggesting that FIP5 and FIP1 function were specific to apical trafficking during lumenogenesis.

Given that FIP1 could compensate for FIP5 in epithelial tissue culture, we asked if Fip1 could do the same *in vivo*. To test this, we performed immunohistochemistry on 6 dpf wild-type, *fip5b^CO40^*, *fip5a^CO35^; fip5b^CO40^*, and *fip5a^CO35^; fip5b^CO40^ mat-* larvae stained for endogenous Fip1 protein. Fip1 staining was mostly absent from wild-type, *fip5b^CO40^* mutant, and *fip5a^CO35^; fip5b^CO40^* zygotic double mutant larvae; however, we observed a significant increase in Fip1 signal in *fip5a^CO35^; fip5b^CO40^ mat-* larvae, especially at the apical cell surface (Figure 4J, K). This suggested that maternal contribution of wild-type *fip5a* may influence Fip1 protein levels to compensate for maternal and zygotic loss of Fip5a together with zygotic loss of Fip5b.

Rab11 specificity for a particular cellular pathway is achieved through interacting with effector proteins, and our work revealed a role for the Rab11 effector paralogs Fip5a and Fip5b in apical cargo delivery and microvilli formation during zebrafish intestinal development. In particular, we observed enlarged Rab7-positive, Rab11-negative organelles in mutants. Normally, there is a homeostasis established between Rab11 recycling from endosomes and maturation from early endosomes to lysosomes (Stenmark, 2009). We propose that without Rab11-Fip5 mediated removal and apical recycling of essential apical cargo, this homeostasis is disrupted such that cargo to be recycled builds up and the maturation process is delayed resulting in engorged Rab7-positive organelles.

One characteristic of MVID is loss of microvilli at the apical cell surface, yet the mechanism behind microvilli phenotypes is still being revealed. Work from intestinal tissue culture and MYO5B mutant mice suggest that disruption of Rab11-mediated recycling of apical membrane proteins and transporters results in failure to maintain apical polarity (Knowles et al., 2014, Vogel et al., 2015, Weis et al., 2016). Our work posits an additional potential explanation in the intermediate filament networks. In polarized epithelia, groups of keratin proteins form polymers at the subapical cell surface just below the apical actin cortex (Apodaca and Gallo, 2013). These actin and intermediate filament networks together comprise the terminal web which is responsible for anchoring the microvilli rootlets into the cell. In *fip5* mutant zebrafish, we observed loss of keratin localization from the apical cell surface to lateral and cytoplasmic regions. It remains unclear how Fip5 regulates apical keratin localization. Because keratins are cytosolic proteins whose assembly and disassembly into networks is mediated by phosphorylation state (Cooper, 2000), one possibility is that Rab11-Fip5 vesicles traffic a keratin kinase or phosphatase to the site of keratin polymerization thereby regulating network assembly. Alternatively, the effect of Fip5 on intermediate filament polymerization could be a more indirect result of general disruption in intracellular trafficking events as we see an accumulation of a number of vesicles and larger organelles in mutant cells. It is interesting to note that in patients with MVID, intractable diarrhea leads to problems with dehydration and electrolyte balance; however, these fish live in an aquatic environment and balance electrolytes through the gills which may mitigate some of these critical problems. In conclusion, our work implicates the Rab11 effectors Fip5 and Fip1 in apical trafficking and microvilli formation during zebrafish intestinal development.

## ABBREVIATIONS

MVID: Microvillus inclusion disease
FIP: Rab11-Family Interacting Protein
dpf: days post-fertilization
MDCK: Madin Darby Canine Kidney

## ACKNOWLEDGEMENTS

We wish to thank Dr. Alexander Blasky for creating the zebrafish *fip5* mutant lines, Dr. Jennifer Bourne for processing zebrafish samples for electron microscopy, and Dr. Michel Bagnat for sharing the hsp:GFP:Rab11a zebrafish line. We are grateful to Drs. Todd Blankenship and Jamie Nichols for critical reading of the manuscript. We are indebted to all members of the Appel lab and the entire zebrafish community at the Anschutz Medical Campus for sharing their protocols, reagents, and knowledge on zebrafish biology. This work was funded by an NSF Graduate Research Fellowship Grant No. DGE-1553798 to C.E.J., Crnic INCLUDE T32 to C.E.J., and NIH 2R01DK064380 to R.P.

## AUTHOR CONTRIBUTIONS

C.E.J. and R.P. performed experiments. C.E.J., B.H.A., and R.P. conceived experiments, wrote the manuscript, and secured funding.

---

**Supplemental Figure 1.**
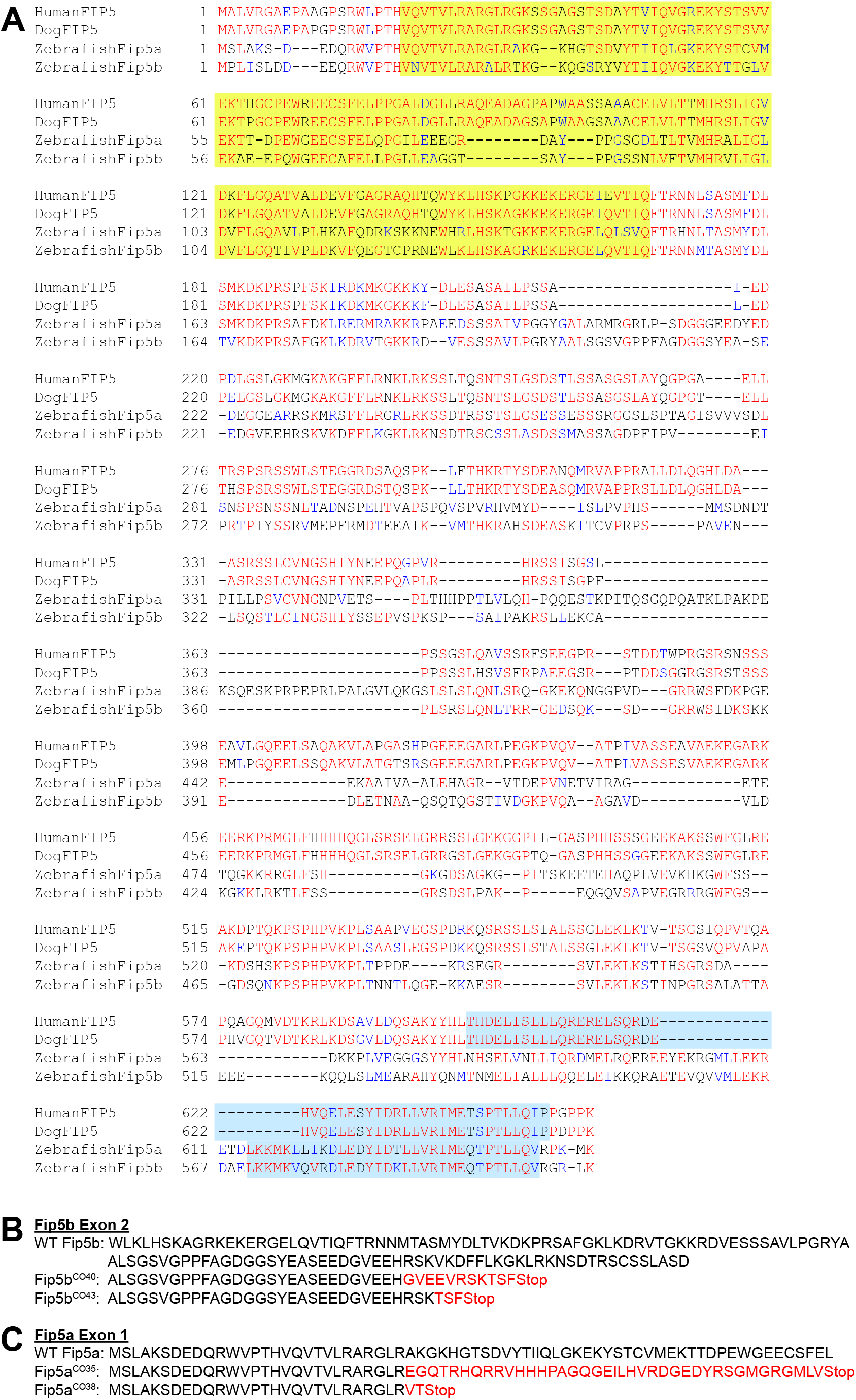
(A) Protein alignments for human FIP5, dog FIP5, and zebrafish paralogs Fip5a and Fip5b. The yellow highlighted region denotes the C2 domain and the blue highlighted region denotes the Rab-binding domain. (B) Fip5b exon 2 sequence in wild-type, *fip5b^CO40^* mutant, and *fip5b^CO43^* mutant alleles. Red amino acids show where mutants differ from wild-type allele. (C) Fip5a exon 1 sequence in wild-type, *fip5a^CO35^* mutant, and *fip5a^CO38^* mutant alleles. Red amino acids show where mutants differ from wild-type allele.

**Supplemental Figure 2.**
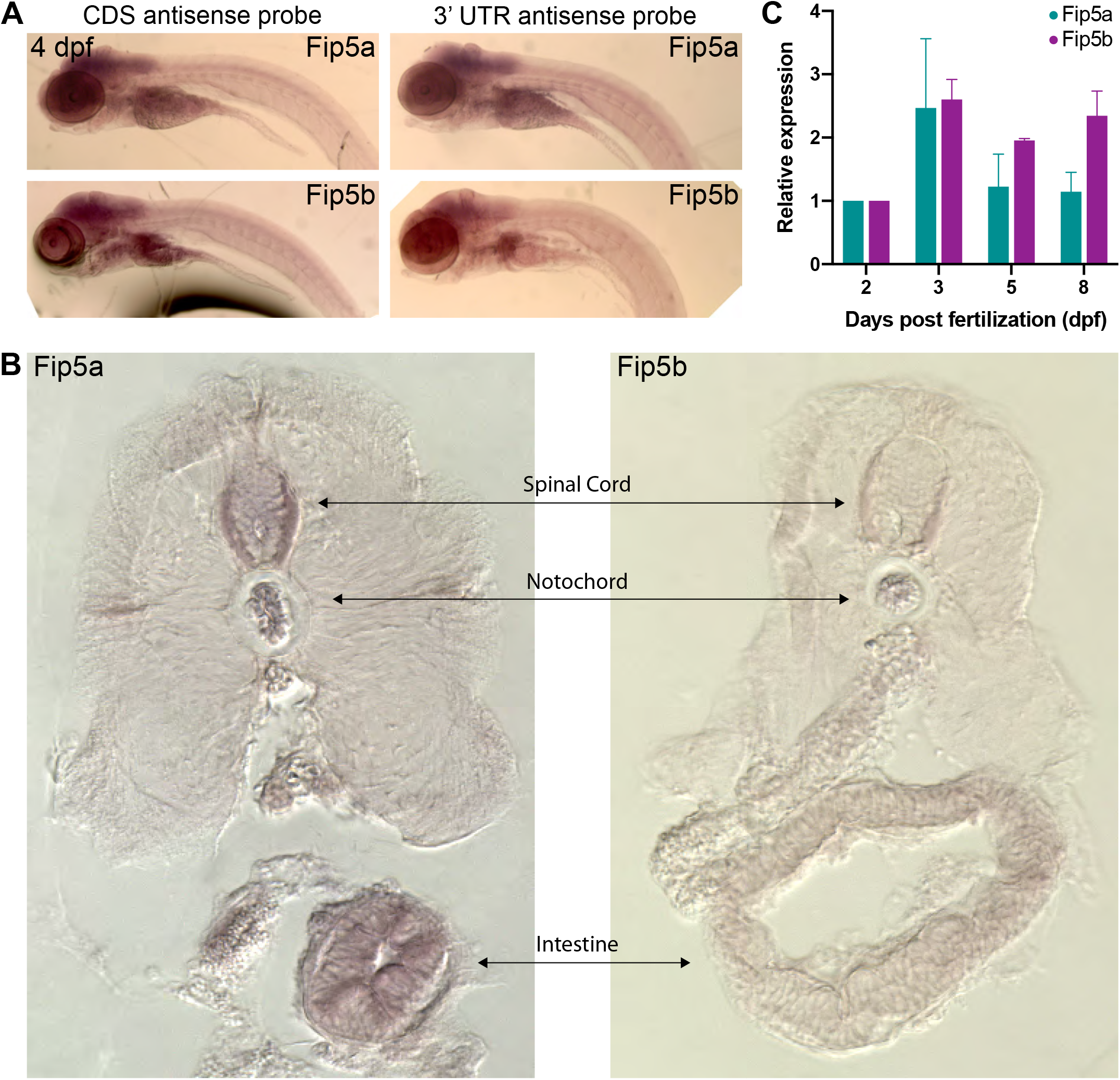
*In situ* hybridization on 4 dpf larvae with antisense probes for the coding sequences of *fip5a* and *fip5b* (left panel) and the 3’ UTR sequences of *fip5a* and *fip5b* (right panel). (B) Representative cross sections of *fip5a* and *fip5b* antisense coding sequence probes. (C) qRT-PCR measuring *fip5a* and *fip5b* transcript levels at 2, 3, 5, and 8 dpf normalized to levels at 2 dpf. All plots show mean with standard error of the mean.

**Supplemental Figure 3.**
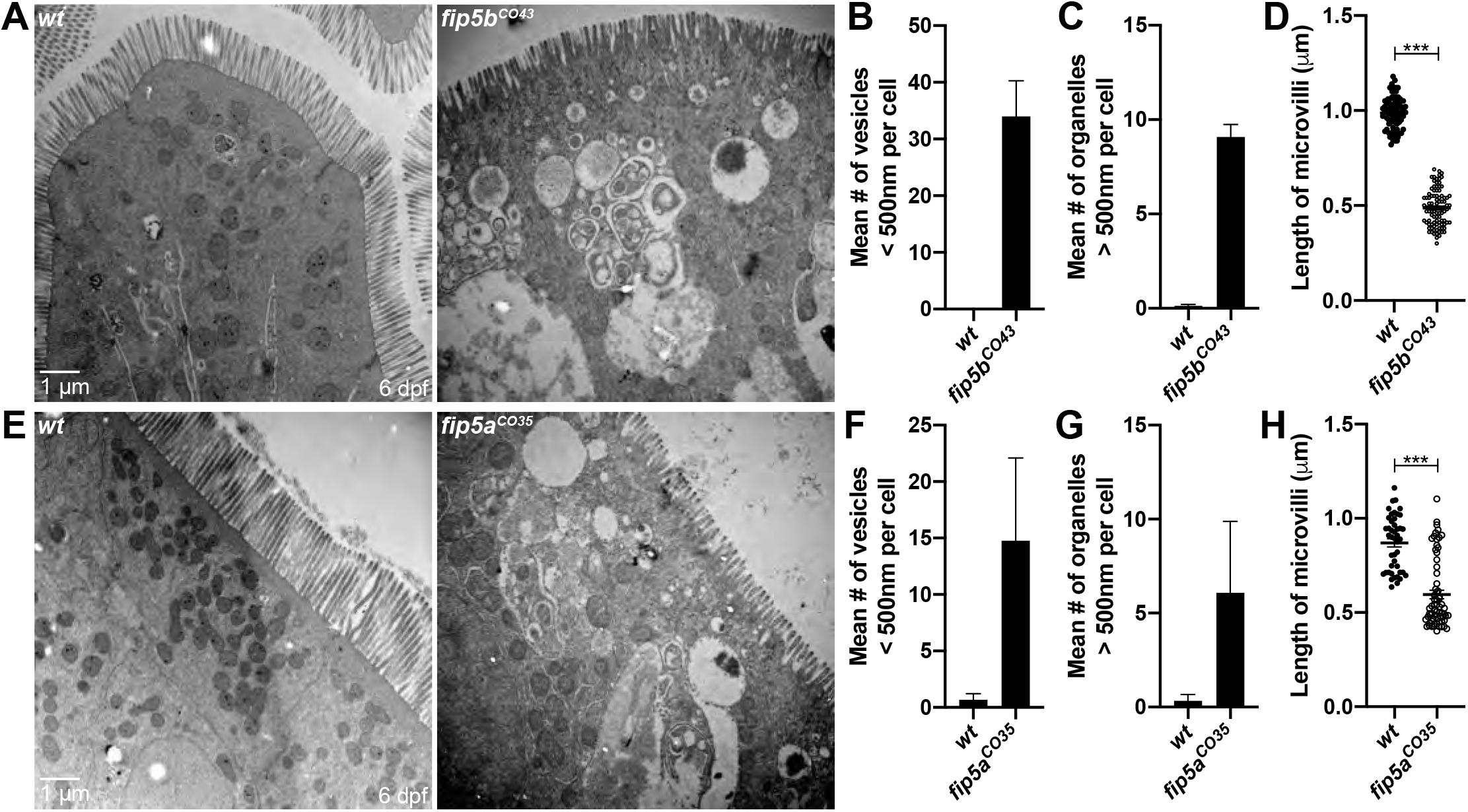
All following images are representative cross sections through midgut region on 6 dpf larvae. Wild-type siblings are used as controls. (A) Electron micrographs showing wild-type and *fip5b^CO43^* mutant larvae. (B) Quantitation of less than 500nm apical vesicles. (C) Quantitation of greater than 500nm organelles. (D) Quantitation of midgut microvilli length. (E) Electron micrographs showing wild-type and *fip5a^CO35^* mutant larvae. (F) Quantitation of less than 500nm apical vesicles. (G) Quantitation of greater than 500nm organelles. (H) Quantitation of midgut microvilli length.

**Supplemental Figure 4.**
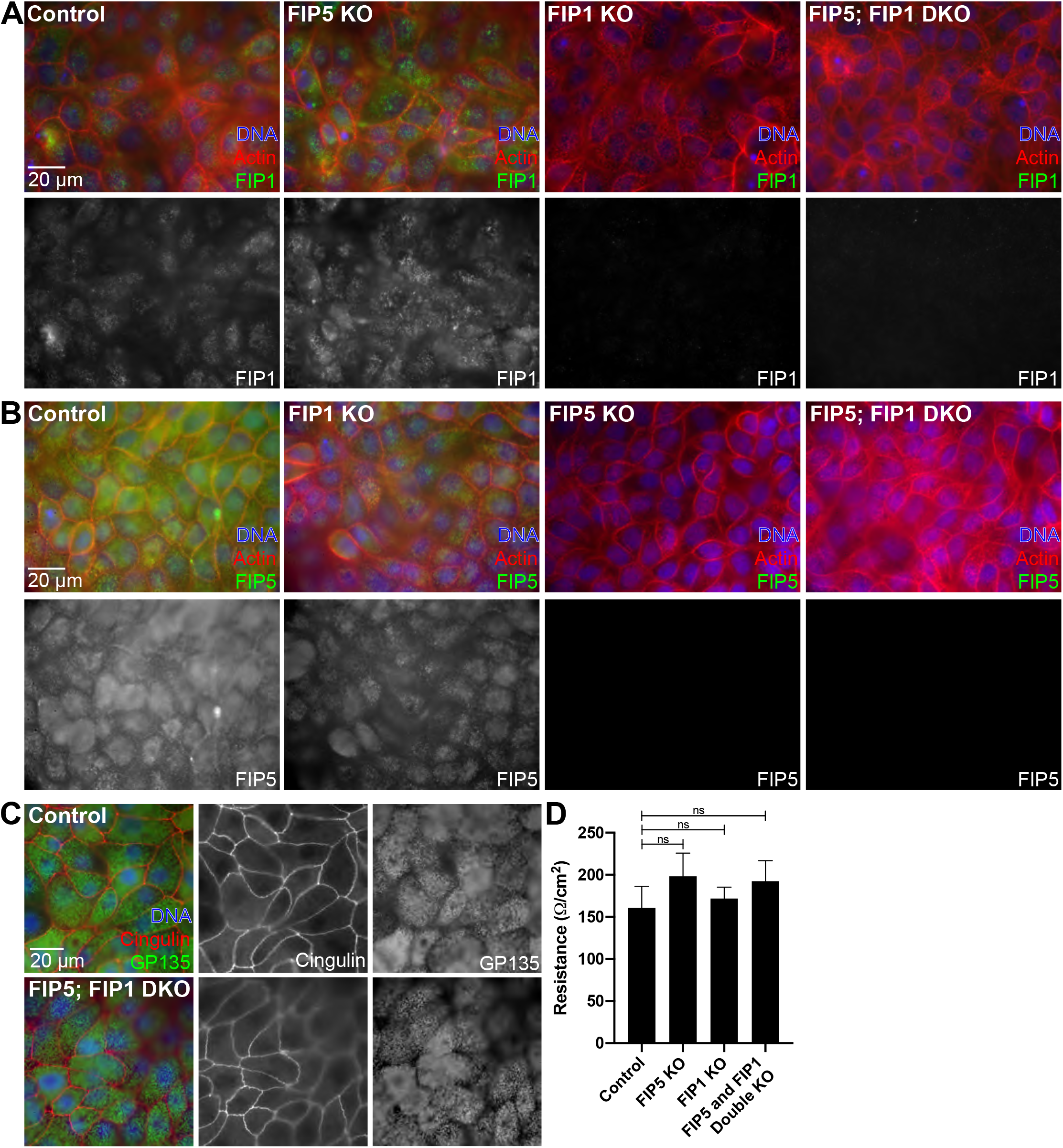
(A) Wild-type, FIP5 KO, FIP1 KO, and FIP5 and FIP1 double KO MDCK cells grown in polarized monolayers and stained for Hoechst (blue), Phalloidin (red) and Fip1 (green). (B) Wild-type, FIP1 KO, FIP5 KO, and FIP5 and FIP1 double KO MDCK cells grown in polarized monolayers and stained for Hoechst (blue), Phalloidin (red) and Fip5 (green). (C) Wild-type and FIP5 and FIP1 double KO MDCK cells grown in polarized monolayers and stained for Hoechst (blue), the tight junction marker Cingulin (red) and the apical membrane marker GP135 (green). (D) Trans-epithelial resistance measurements on wild-type, FIP5 KO, FIP1 KO, and FIP5 and FIP1 double KO MDCK cells grown in polarized monolayers.

## MATERIALS AND METHODS

### Zebrafish husbandry

All stocks unless otherwise specified were maintained in a heterozygous state and kept according to Standard Operating Procedure defined in “The Zebrafish Book” (M. Westerfield, Inst. of Neuroscience, Univ. of Oregon).

### qRT-PCR

RNA extraction from larvae was performed with TRIzol reagent (Invitrogen) followed by cDNA synthesis with iScript cDNA Synthesis Kit (BioRad). SYBR Green PCR Master Mix (Applied Biosystems) was used for qPCR. All reactions were performed in technical triplicate and a minimum of three biological replicates were performed. Primer sequences are listed in Table 2.

### Protein alignments

Fip5 protein alignments were generated using T-Coffee and Boxshade 3.2. The following protein accession numbers from NCBI were used for alignments: Human NP_056285; Dog XP_003639656 (isoform X5); Zebrafish Fip5a XP_009305489 (isoform X2); Zebrafish Fip5b XP_017214658 (rab11 family-interacting protein 5-like isoform X2).

### Zebrafish Immunohistochemistry

Larvae were placed in 1-2% Tricaine for 10 minutes or until they were unresponsive to touch then decapitated immediately posterior to the otic vesicle using a scalpel. The larva body was placed in fix solution (4% paraformaldehyde, 4% sucrose, 0.15 mM CaCl_2_, pH 7.3) at 4°C overnight, whereas the head was placed in lysis buffer and genotyped (see genotyping). The fixed larvae were then embedded in a melted agar solution (1.5% agar, 5% sucrose in water), and after the blocks hardened, they were trimmed and immersed in 30% sucrose in water solution overnight at 4°C. Blocks were then dried with a chemwipe, frozen on dry ice for ~15 minutes, then stored at −80°C until ready to section. Blocks were mounted in OCT and 20um sections cut using a Leica CM 1950 cryostat microtome. Sections were placed on FisherBrand charged slides (Cat # 12-550-15) and rehydrated in PBS for 30 minutes. Excess liquid was dried, and then a wax pen was used to draw around the edge of the slide. Slides were then blocked with 2% BSA and 5% donkey serum (ThermoFisher Cat # NC9624464) in PBS for 1 hour, then incubated in primary antibody (see antibodies in Table 1) diluted in block at room temperature for 2-3 hours. Slides were then washed 4x 15 minutes each with PBS and incubated in secondary antibody (see antibodies in Table 1) diluted in block for 1-2 hours at room temperature. Slides were again washed 4x 15 minutes each with PBS, adding Hoescht (ThermoFisher Cat # 33342 at 1:500) to the second to last wash. Slides were then dried, mounted in Vectashield (Vector Laboratories Cat # H-100), and sealed with nail polish.

**Table 1:** Antibodies

| Name | Supplier/Cat # | Dilution |
| --- | --- | --- |
| Pan cytokeratin AE1/AE3 | Abcam/ab27988 | 1:50 |
| Rab7 | Abcam/ ab50533 | 1:100 |
| Rab11 | Life Technologies/ 715300 | 1:100 |
| GP135 | DSHB/ 3F2/D8 | 1:100 |
| Cingulin | Prekeris Lab | 1:100 |
| FIP1 | Prekeris Lab | 1:200 |
| FIP5 | Prekeris Lab | 1:100 |
| Alexa 488 Anti-Rabbit secondary | Jackson ImmunoResearch/711-545-152 | 1:100 |
| Alexa 488 Anti-Mouse secondary | Jackson ImmunoResearch/715-545-150 | 1:100 |
| Alexa-568 Phalloidin | Invitrogen/A12380 | 1:100 |

**Table 2:**
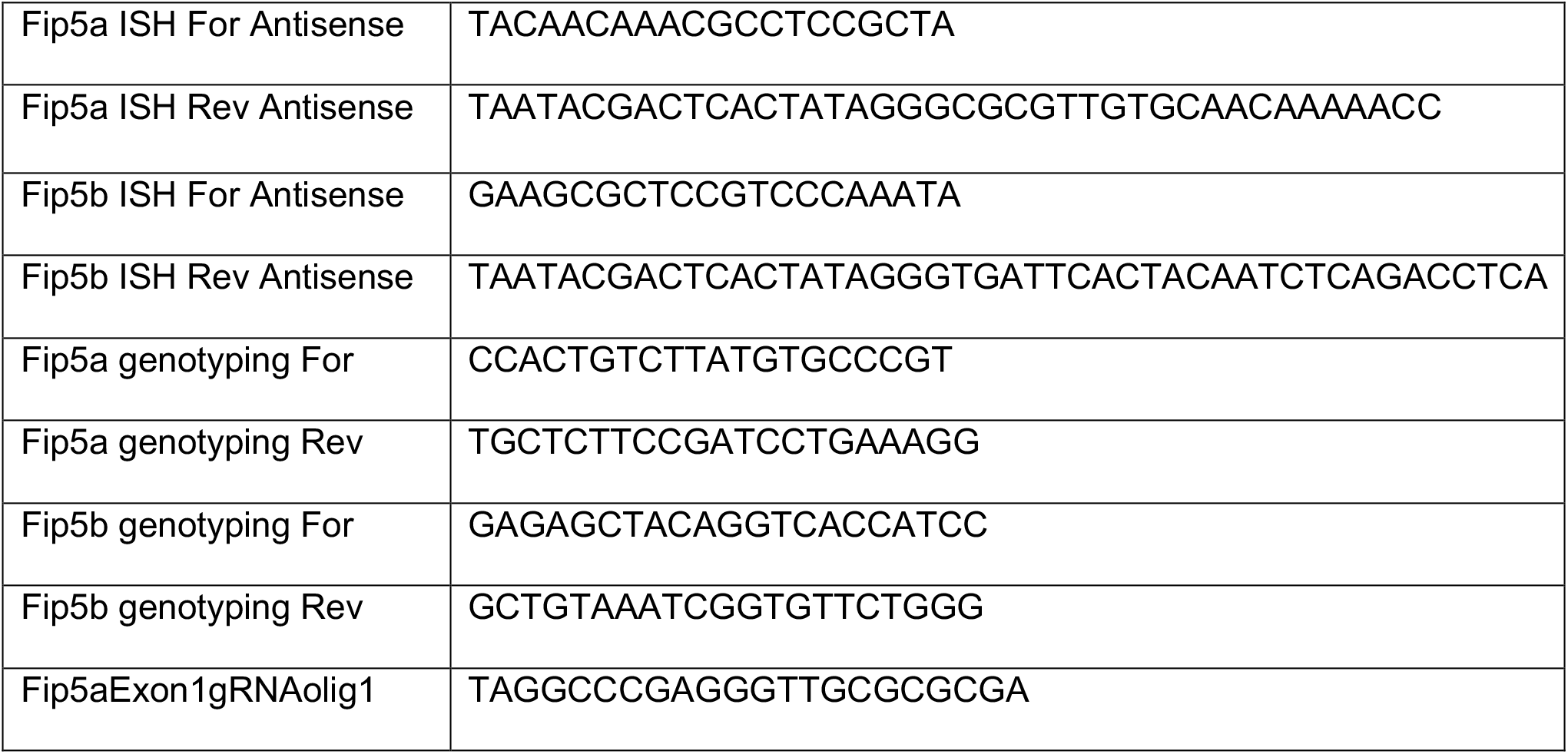
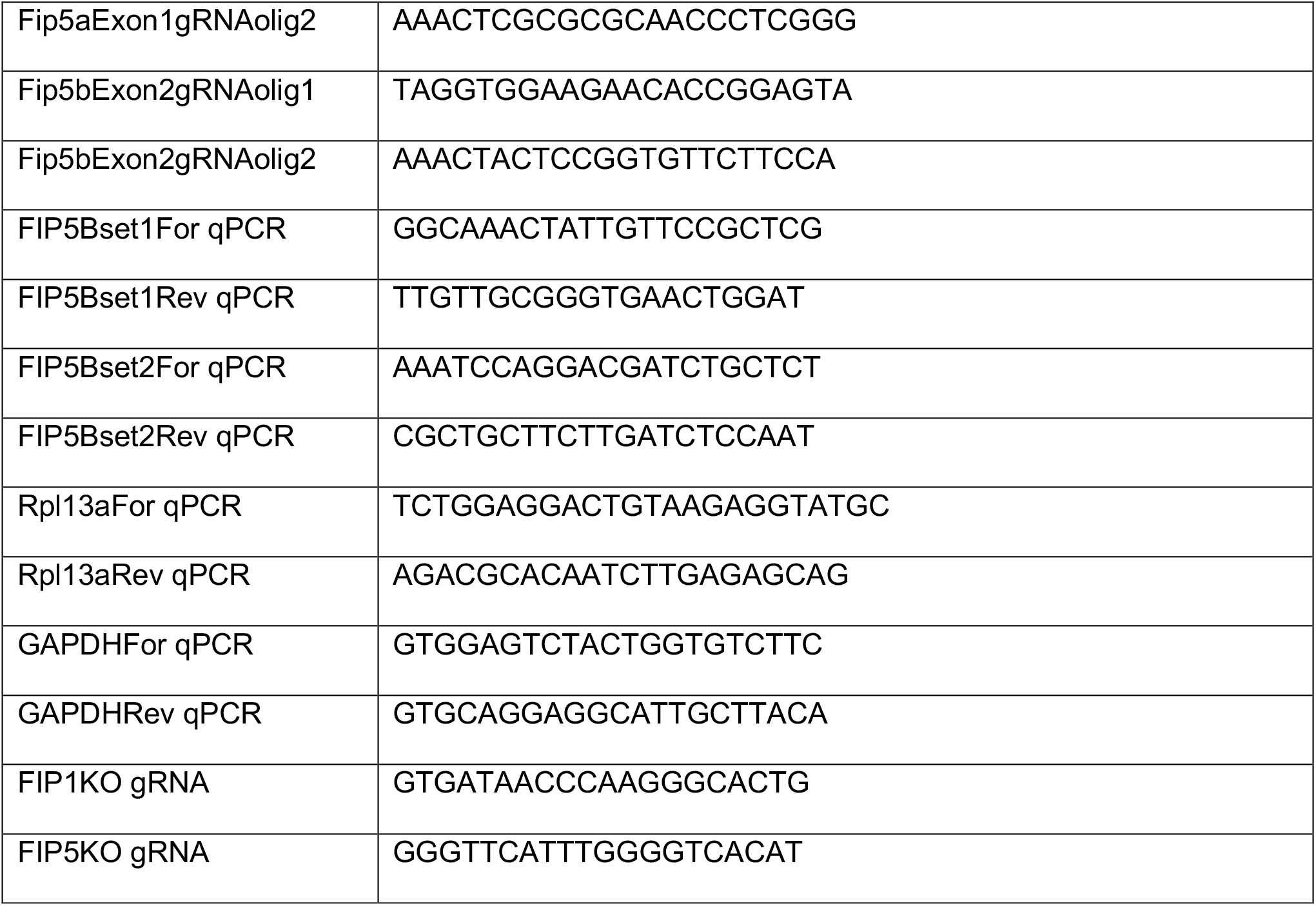
Primer sequences. Primers were designed using the NCBI/Primer-BLAST tool.

### Widefield Microscopy and Image Analysis

All slides of fixed fish sections were imaged with an inverted Axiovert 200M microscope (Carl Zeiss) with a 63x oil immersion lens and QE charge-coupled device camera (Sensicam). Images were acquired using Slidebook 6.0 (Intelligent Imaging Innovations) software. Images were processed using a combination of Slidebook 6.0 (Intelligent Imaging Innovations) software, Fiji (PMID 22743772), and Adobe Photoshop. Figures were made in Adobe Illustrator. A minimum of three biological replicates were performed for each experiment and quantitation was performed unblinded.

### Genotyping Zebrafish

Fish tissue was isolated from a fin clip for adult fish or from the heads for larvae. Fish tissue was placed in lysis buffer (10 mM Tris pH 8.3, 50 mM KCl, 1.5 mM MgCl_2_, 0.3% Tween-20, 0.3% NP-40 in water) with 2% Proteinase K (Invitrogen Cat # 25530049). Lysis reactions were incubated at 55°C for 4 hours, then 95°C for 20 minutes to inactivate Proteinase K. A PCR/Restriction Enzyme-based assay was used to genotype *fip5a* and *fip5b* mutant fish lines. For *fip5a*, a 400bp region of genomic DNA surrounding the CRISPR target site was amplified by PCR. The PCR product was then digested with BssHII for 1 hour at 50°C, and the resulting product was run on a 2% agarose gel. For *fip5b*, a similar schematic was used with the BsaWI or AgeI restriction enzyme depending on the allele. PCR primer sequences are listed in Table 2. Genotyping was performed prior to experiments and only wild-type and homozygous mutant larvae were selected for analysis.

### CRISPR/Cas9 in Zebrafish

All primer sequences are listed in Table 2. Guide RNA (gRNA) oligos were designed using ZiFIT Targeter Software for CRISPR/Cas9 Nucleases (Sander et al., 2010, Sander et al., 2007). The gRNA target sites were then blasted (NCBI Blastn) against the zebrafish genome to look for potential off target sites. gRNA oligos were annealed and phosphorylated, then cloned into the *pDR274* vector (Addgene) using the BsaI-HF restriction site. Positive clones were sequenced to confirm correct insertion. The gRNA-containing vector was linearized using DraI and purified by ethanol precipitation. The gRNA sequence was then transcribed to RNA using T7 polymerase and purified by phenol choloroform extraction. Cas9 mRNA was synthesized from the pT3TS vector (Addgene) using mMESSAGE mMACHINE T3 Transcription Kit (ThermoFisher Cat# AM1348) and purified using phenol choloroform extraction. The injection mix was prepared as follows: 0.2 M potassium chloride, 0.15 ng uL^−1^ Cas9 mRNA, 70ng uL^−1^ gRNA, and 10% phenol red in DEPC water. Embryos were injected with 1-3 nL of the injection mixture at the 1-cell stage. Founder fish were determined using T7E1 analysis (NEB), and positive hits were sequenced to determine exact mutation. Founders from two different gRNA injection experiments containing different mutant alleles for *fip5a* and *fip5b* were selected and outcrossed for at least two generations before performing experiments.

### Transmission Electron Microscopy

Larvae were placed in 1-2% Tricaine for 10 minutes or until they were unresponsive to touch then decapitated immediately posterior to the otic vesicle. The larva body was placed in EM fix solution (0.1M sodium cacodylate, 4% paraformaldehyde, 4% glutaraldehyde, in PBS) at 4°C overnight, whereas the head was placed in lysis buffer and genotyped. The body was processed for EM by washing in 0.1M sodium cacodylate, then incubating tissue in 500uL of 1:1 osmium tetroxide to 0.1M sodium cacodylate for 2 hours. Tissue was washed with double distilled water, then incubated in 500uL 1:1 osmium tetroxide to imidazole (0.35g imidazole in 25mL sodium cacodylate pH to 7.4) for 30 minutes. Larvae turned brown at this point. Larvae were washed again in double distilled water then an ethanol dehydration series was performed (50% / 75% / 100%). Larvae were then incubated in 1:1 Epon to ethanol for 1 hour, then 2:1 Epon to ethanol overnight. The following day, larvae were embedded in 100% Epon, which was replaced with fresh 100% Epon, and then baked for 2 days. Larvae were cut in half where the body narrows (see schematic in Figure 1C), and then 65 nm thick sections were cut and collected on formvar-coated copper slot grids. Sections cut in the anterior direction were designated the intestinal bulb region and in the posterior direction the midgut. Sections were imaged on a FEI Tecnai G2 Biotwin Transmission Electron Microscope, run at 80 kV with a side-mount AMT XR80S-B digital camera. For TEM quantitation, a minimum of three biological replicates were used for each experiment and images were blinded prior to analysis.

### RNA In Situ Hybridization

Sense and antisense RNA probes were designed against both the coding sequence and 3’ UTR region of zebrafish *fip5a* and *fip5b* genes. A PCR-based method with T7 sites at the end of the primers was used to amplify the probe DNA sequence from 8 day post fertilization wild-type fish cDNA (see Table 2).

The RNA probes were transcribed with the T7 polymerase and labeled using the DIG RNA labeling kit (Roche Cat # 11175025910). After the labeling reaction was complete, the probes were mixed with 1 μl 0.5M EDTA to stop the reaction, then 2 μl 5M lithium chloride, and 75 μl cold ethanol were added for purification by ethanol precipitation and the probe was resuspended in DEPC water. The RNA probe was checked by agarose gel electrophoresis, then mixed with an equal volume of formamide and stored at - 80°C.

RNA in situ hybridization assays were conducted based on a modified version of a previously published protocol described by Hauptmann and Gerster (Hauptmann and Gerster, 2000). Larvae were fixed in 4% paraformaldehyde in DEPC PBS overnight at 4°C. Larvae were stored in MeOH at −20°C until use, when they were washed twice for five mins in DEPC-PBSTw (0.5% Tween-20 in PBS made with DEPC water). Pigmentation was bleached in a hydrogen peroxide solution (3% H_2_O_2_, 0.5% KOH in DEPC water) until larvae eyes turned brown (15-30 minutes). Larvae were then washed twice for 5 mins in DEPC-PBSTw, fixed for 20 minutes at room temperature in 4% PFA in DEPC-PBS, and washed again twice for 5 mins in DEPC-PBSTw. The larvae were digested with 0.1 mg/mL Proteinase K (Invitrogen Cat # 25530049) in DEPC-PBS for 17 minutes to permeabilize the larvae, then washed twice for 5 mins each wash in DEPC-PBSTw, followed by fixation for 20 minutes in 4% PFA, and again washed twice for 5 mins in DEPC-PBSTw. Larvae were incubated in 500 μL Hybridization Media Block solution (50% formamide, 5x Saline-Sodium Citrate Buffer, 10 μL/mL tRNA, 50 mg/mL heparin, 0.01M citric acid, and 0.5% Tween-20 in DEPC H_2_O) for 1 hour at 70°C. The block was replaced with Hybridization Media containing 200 ng of the appropriate RNA probe, and the larvae were incubated at 70°C overnight. The following day, a series of progressive washes were performed for 10 mins each wash at 70°C: 200 μL 100% HM without probe, 300 μL 66% HM / 33% 2x Saline-Sodium Citrate Buffer (SSC; Cellgro, Mediatech, Inc., Manassas, VA), 300 μL 33% HM / 66% 2x SSC, 1 mL 2x SSC, 1 mL 0.2x SSC, 1 mL 0.1 x SSC (this wash was performed twice), 1 mL DEPC-PBSTw. Another 10 min wash with DEPC-PBSTw was performed at room temperature, followed by an hour-long antibody block (2% sheep serum and 2 mg/mL BSA in DEPC-PBSTw). Anti-Digoxigenin-AP Fab fragments (Roche Applied Science, Indianapolis, IN) was incubated in antibody block overnight at 4°C. Finally, four 15 min washes were conducted at room temperature in DEPC-PBSTw. Larvae were incubated in staining solution (0.1M Tris, pH 9.5, 0.25M MgCl_2_, 0.1M NaCl, 0.5% Tween-20) for 15 minutes at room temperature. Thereafter, the larvae were moved to a staining dish, covered with 500 μL precipitating BM Purple AP Substrate (Roche Applied Science, Indianapolis, IN), and incubated at 37°C for 8 hours until staining was visible. The larvae were then washed twice for 5 mins in PBS and imaged or processed for sectioning immediately.

### Cell culture and immunohistochemistry

MDCK II cells (ATCC) were cultured in DMEM with 10% FBS and penicillin/streptomycin. For polarized MDCK experiments, cells were plated on collagen type I-coated Transwell filters (Corning 3460) to reach confluency in 24 hours. Cells were then grown for three more days before transepithelial resistance measurements or fixation. Cells were fixed with 4% paraformaldehyde for 20 minutes at room temperature Cells were blocked for 1-2 hours in block buffer (PBS, 0.1% Triton X-100, 10% normal donkey serum). Primary antibodies were diluted in block buffer and incubated overnight at room temperature. Cells were washed with PBSTx before adding secondary antibodies for 1-2 hours at room temperature. Cells were washed again before mounting in VectaShield and sealing with nail polish or mounting in Prolong Gold. Coverslips used for all experiments were #1.5 thickness.

### Trans-epithelial resistance measurements

MDCK cells were grown on collagen-coated transwell filters (see Cell Culture section) and resistance measurements were taken four days after plating with a Millicell ERS-2 Voltohmmeter (Millipore). Three measurements per well, one for each space between the plastic prongs of the filter holder, were averaged and subtracted from the average of the blank well containing a collagen-coated filter without any cells.

### Generating MDCK and RPE-1 CRISPR knockout lines

MDCK cells stably expressing Tet-inducible Cas9 (Dharmacon Edit-R inducible lentiviral Cas9 nuclease) were grown in a 12-well dish to about 75% confluency before treatment with doxycycline at a final concentration of 1ug/mL for 24 hours to induce Cas9 expression. Cells were then transfected with crRNA:tracrRNA mix as described for DharmaFECT Duo co-transfection protocol (Horizon Discovery Cat# T-2010-xx). Transfected cells were incubated for 24 hours before trypsinizing and plating for individual clones. Individual clones were screened through genotyping PCR and sanger sequencing. All CRISPR gRNAs are listed in Table 2.

